# Caloric restriction increases levels of taurine in the intestine and stimulates taurine uptake by conjugation to glutathione

**DOI:** 10.1101/2020.07.23.217612

**Authors:** András Gregor, Marc Pignitter, Christine Fahrngruber, Sebastian Bayer, Veronika Somoza, Jürgen König, Kalina Duszka

**Affiliations:** Department of Nutritional Sciences, University of Vienna, Vienna, Austria; Department of Physiological Chemistry, University of Vienna, Vienna, Austria; Leibniz-Institut for Food Systems Biology, Technical University of Munich, Germany

**Keywords:** caloric restriction, bile acids, taurine, glutathione, intestine

## Abstract

Our previous study indicated increased levels of taurine-conjugated bile acids in the intestine content of caloric restriction (CR) mice. In the current project, we found increased levels of free taurine and taurine conjugates, including glutathione (GSH)-taurine, in CR compared to *ad libitum* fed animals in the mucosa along the intestine while there was no impact on taurine and its conjugates in the liver. The levels of free GSH were decreased in the intestine of CR compared to *ad libitum* fed mice. However, the levels of oxidized GSH were not affected and were complemented by the lack of changes in the antioxidative parameters. Glutathione-S transferases (GST) enzymatic activity was increased as was the expression of GST genes along the GI tract of CR mice. In CR intestine addition of GSH to taurine solution enhanced taurine uptake. Accordingly, the expression of taurine transporter (TauT) was increased in the ileum of CR mice.

## Introduction

Caloric restriction (CR) is one of the primary intervention tools applied for weight loss and health maintenance, showing remarkable health benefits. CR lowers the incidence of multiple diseases and diminishes the rate of age-specific mortality, resulting in extended lifespan (Speakman and Mitchell, 2011, Weindruch et al., 1986, Anderson et al., 2009b, Fontana et al., 2010, Mattison et al., 2017). A reduction of oxidative stress is claimed to be among the factors contributing to the beneficial outcomes of CR (Merry, 2002, Sanz et al., 2005, Yu, 1996). Reduced glutathione (GSH) is one of the most important non-enzymatic antioxidants which serves as a scavenger of free radicals, aids in the reduction of H_2_O_2_, and takes part in detoxification. Together with glutathione peroxidase (GPx) and glutathione S-transferases (GSTs), GSH forms the glutathione system, which is abundant in the GI tract (Siegers et al., 1988, Hoensch et al., 2002). GSTs function as a family of enzymes, which catalyze the conjugation of GSH to different electrophilic substrates, thereby producing water-soluble compounds that are further directed to excretion via urine and bile (Hinchman and Ballatori, 1994). Bile is produced by the liver and mainly consists of water, bilirubin, and bile acids (BA). Upon secretion to the small intestines, BAs enhance lipid uptake. Once formed, BAs are conjugated to glycine or taurine. Taurine is one of the most abundant free amino acids in the body (Schuller-Levis and Park, 2003). It has anti-inflammatory (Kim and Cha, 2014, Marcinkiewicz et al., 2005, Niu et al., 2018, Chupel et al., 2018), antioxidative (Silva et al., 2011, Niu et al., 2018, Thirupathi et al., 2018, Bucolo et al., 2018, Lee et al., 2019), and osmoprotective (Bucolo et al., 2018, Trachtman et al., 1988) properties. In the intestine of an immunosuppressive mouse model, taurine increases the number of some immune cells and total cells in Payer’s patches (Fang et al., 2019), it reduces the growth of harmful bacteria, increases the production of short-chain fatty acids (Yu et al., 2016), and attenuates induced colitis (Shimizu et al., 2009, Zhao et al., 2008). In human intestinal epithelial Caco-2 cells, taurine stimulates the expression of anti-inflammatory factors (Gondo et al., 2012).

As reported previously (Duszka et al., 2018), we measured increased levels of the taurine-conjugated BA taurocholic acid (TCA) and tauroursodeoxycholic acid (TUDCA) in the intestinal content of mice submitted to CR compared to *ad libitum* feeding. In the current study, we explored the consequences of CR-increased levels of intestinal taurine-conjugated BA. We hypothesized that CR-associated elevated levels of taurine-conjugated BA are accompanied by increased levels of free taurine steaming from deconjugation in the intestine. Since taurine is a bioactive compound we wanted to investigate the consequences of this regulation.

## Results

As we previously published, CR is associated with increased levels of taurine-conjugated BA in the intestine (Duszka et al., 2018). To further explore this phenomenon, we studied the impact of CR on intestinal taurine levels. In order to mirror previous experimental conditions, male C57Bl/6 mice were submitted to 14 days caloric restriction receiving 75% of the amount of standard chow that the animals would voluntarily consume. The levels of taurine were measured in the mucosa along the small intestine. We detected increased levels of free taurine in the duodenum of CR compared to *ad libitum* fed mice (Figure 1A). To verify how taurine could be utilized in the intestine, compounds containing taurine were analyzed. Multiple conjugates of taurine were detected and, accordingly, their levels were increased in the mucosa of CR animals (conjugate with molecular mass 249 is presented) (Figure 1B). Among the compounds detected, we identified a GSH-taurine conjugate. The conjugate showed an increased concentration in intestinal mucosa samples of CR compared to *ad libitum* fed mice. However, in the duodenum, this difference was not statistically significant (Figure 1C). The levels of free taurine, taurine conjugates, and GSH-taurine were statistically significantly elevated in the jejunum (Figure 1D-F), and the ileum of CR animals (Figure 1G-I). The differences in the amounts of the metabolites were more pronounced in jejunum and ileum compared to the duodenum. The level of one of the taurine metabolites, taurine chloramine (TauCl), a compound with strong anti-inflammatory properties, was also increased in the ileum (Suppl Figure 1A). We further verified if CR affects also liver in a similar way, but there was no difference in free taurine (Figure 2A), conjugated taurine (Figure 2B), or GSH-taurine conjugates (Figure 2C) levels between CR and *ad libitum* mice. Importantly, the expression of genes associated with BA synthesis (*Cyp7a1*), transport (*Ntcp*), and cysteine metabolism (*Cdo*) was upregulated in CR liver while that of the taurine transporter gene (*Taut*) was not affected by the diet (Figure 2A).

**Figure 1.**
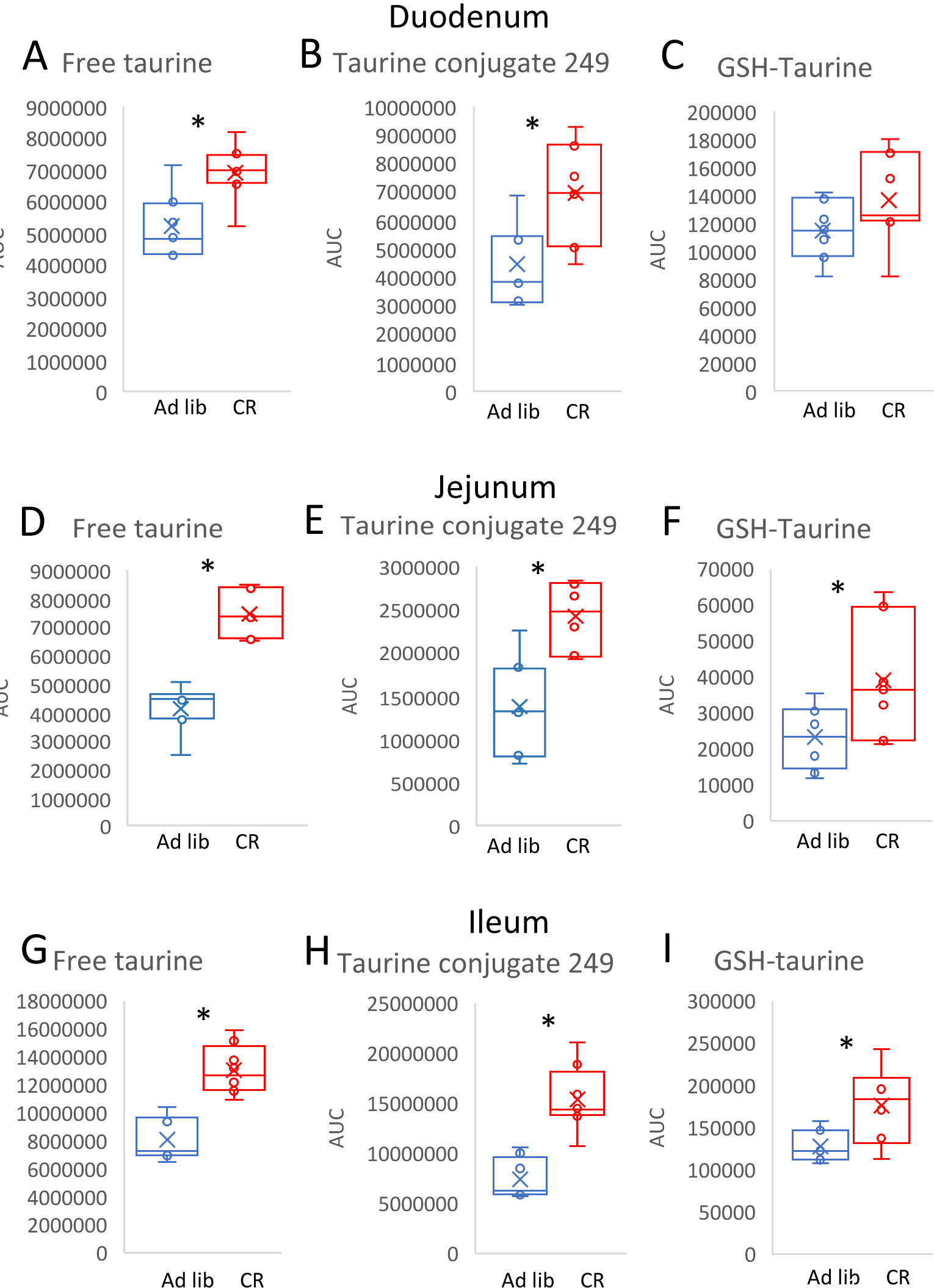
CR increases the levels of free taurine and its conjugates along the intestinal mucosa. The levels of free taurine (A, D, G, J), taurine conjugate with m/z 249 (B, E, H) and GSH-taurine conjugate (C, F, I) were measured in the duodenum (A-C), jejunum (D-F), and ileum (G-I). Statistical significance was assessed using a two-tailed Student’s t-test; *p<0.05; n=6-8. Error bars stand for ±SEM.

**Figure 2.**
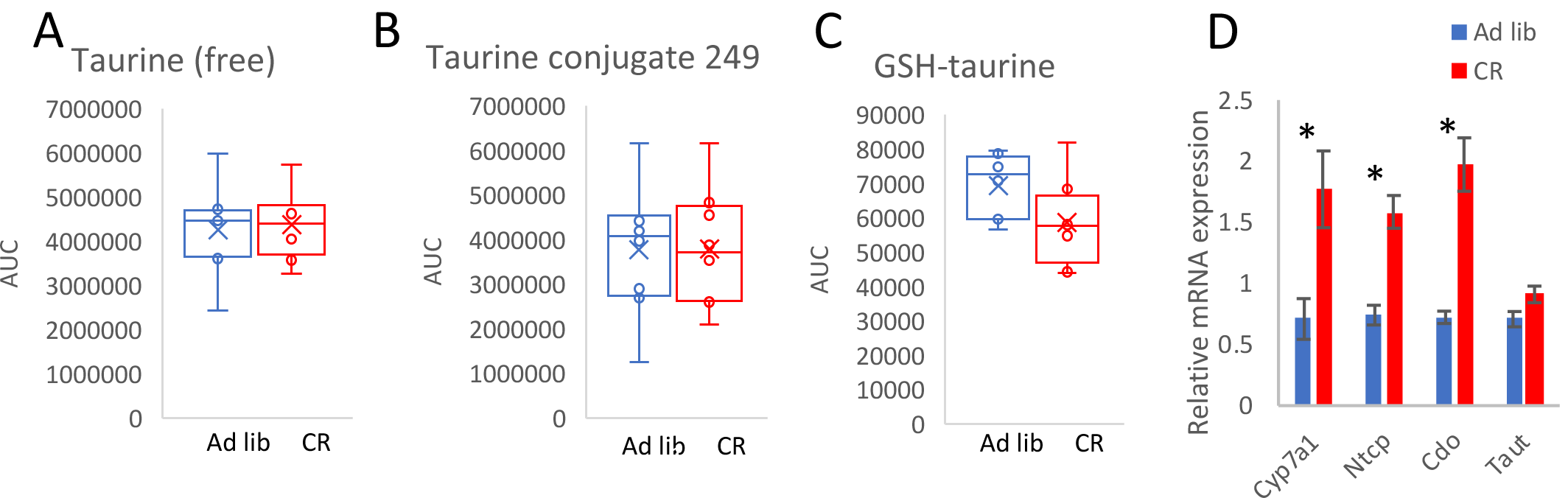
CR does not affect the levels of taurine and its conjugates in the liver but regulates BA- and cysteine-related gene expression. The levels of free taurine (A), taurine conjugate with m/z 249 (B), and GSH-taurine conjugate (C) were measured in the liver. Gene expression was quantified in the liver of CR and *ad libitum* fed mice (A). Two-tailed Student’s t-test was applied to assess statistical differences between the groups; *p<0.05. Bars indicate the mean of eight to nine biological replicates ±SEM.

Since one of the identified taurine conjugates contained GSH, we verified the levels of reduced (GSH) and oxidized (GSSG) glutathione in the jejunum mucosa. The levels of GSH were significantly decreased in the CR compared to *ad libitum* fed mice (Figure 3A), while GSSG concentration was not affected by CR (Figure 3B). To validate whether the rate of *de novo* synthesis or utilization of GSH is modified during CR, expression of genes connected with GSH synthesis (nuclear factor erythroid 2-related factor 2 (*Nrf2*), gamma-glutamyl transpeptidase (*Ggt*), glutamate-cysteine ligase (*Gcl*), and glutathione synthase (*Gs*)) and antioxidant activity (glutathione reductases (*Grx1, Grx2*) and glutathione peroxidases (*GPx1, GPx2*), as well as activity of several of the factors, were measured in the mucosa of the jejunum.

**Figure 3.**
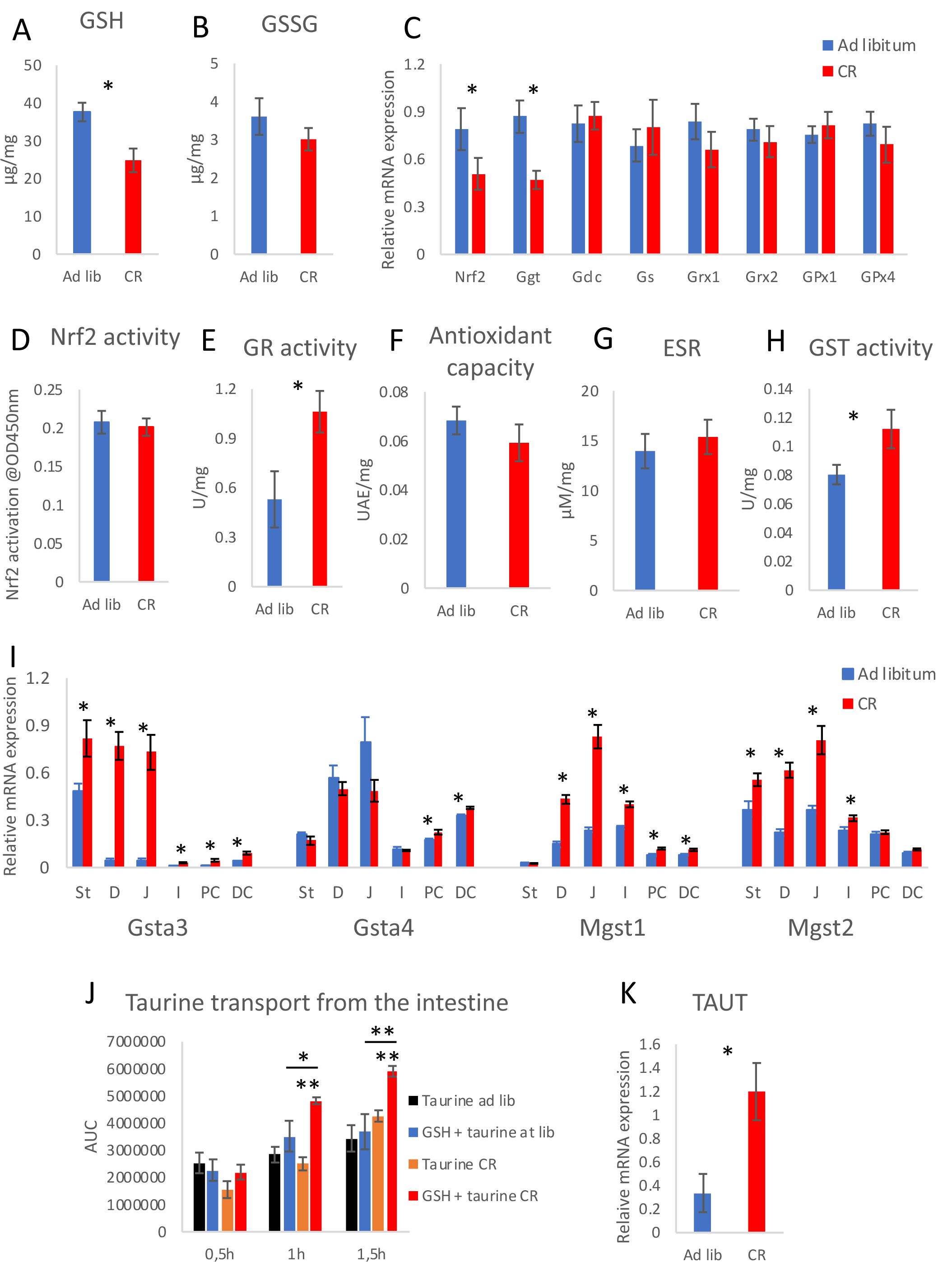
Caloric restriction (CR) impacts glutathione (GSH) and glutathione-S transferase (GST) in the jejunum mucosa. The levels of reduced GSH (A) and oxidized GSH (GSSG) (B) were measured in the jejunum mucosa using commercial assays. Gene expression was measured in the mucosa of the jejunum of CR and *ad libitum* mice (C). The activity of nuclear factor erythroid 2-related factor 2 (Nrf2) (D) and glutathione reductase (GR) (E) as well as antioxidant capacity (F), were measured in the mucosa of the jejunum utilizing commercial assays. Electron spin resonance (ESR) was applied to assess the levels of reactive oxygen species in the mucosa of the jejunum (G). GST activity was measured in the jejunum mucosa using commercial assays (H). Gene expression of the different GST subtypes was measured in the mucosa of six parts of the GI tract (St-stomach, D-duodenum, J-jejunum, I-ileum, PC-proximal colon, DC-distal colon) of CR and *ad libitum* fed mice (I). Taurine uptake was assessed using intestine sacs infused with solution containing taurine or taurine and GSH (J). The expression of *TauT* mRNA was measured in the mucosa of the ileum (K). Two-tailed Student’s t-test was applied to assess statistical differences between the groups; *p<0.05, **p<0.001. Error bars indicate ±SEM. Figures in panels A-I and K represent the mean of eight to nine biological replicates while in panel J mean of five to seven replicates.

The mRNA level of *Nrf2* was downregulated in the CR mice (Figure 3C), although its transcriptional activity was not changed (Figure 1D). Also, the expression of *Ggt* was downregulated (Figure 3C). The expression of other genes connected with GSH synthesis and its anti-oxidative role was not affected (Figure 3C). The activity of GR which is responsible for the conversion of GSSG to GSH was increased in the CR animals (Figure 3E), whereas the activity of GPx was not significantly affected and showed high variability (Suppl Figure 1B). To verify the functional outcome of the anti-oxidative role of GSH in the CR intestine, the total antioxidant capacity was assessed (Figure 3F) and the levels of reactive oxygen species were measured by means of electron spin resonance (ESR) (Figure 3G) in the intestinal mucosa. Both of the parameters were not affected. Therefore, we decided to focus on the conjugating function of GSH and found increased activity of GST in the intestine of CR compared to *ad libitum* fed mice (Figure 3H). We also measured gene expression in six distinct parts of the GI tract (stomach, duodenum, jejunum, ileum, proximal colon, and distal colon) (Figure 3I). Fittingly with the enzymatic activity results, the expression of *Gst* genes was consistently increased in mucosa of the different parts of the GI tract. However, the range of upregulation varied between different forms of *Gst*. To verify the functionality of the occurrence of GSH-taurine conjugates, we incubated *ex vivo* intestinal sacs of CR and *ad libitum* fed mice which were infused with taurine or mix of taurine and GSH solutions to measure uptake of taurine. The levels of taurine in the surrounding medium were measured at three time points over 1,5h of incubation. The uptake of taurine did not change between CR and *ad libitum* fed mice when sacs were infused with a taurine solution (Figure 3J). Similarly, a solution containing taurine and GSH did not impact taurine uptake in *ad libitum* fed mice. However, incubation of the intestine of CR mice with taurine and GSH solution strongly increased the concentration of taurine in the surrounding medium. Correspondingly, the mRNA expression of *Taut* was higher in ileum mucosa of CR compared to *ad libitum* fed mice (Figure 3K).

## Discussion

To contribute to building the full picture of the molecular mechanism of intestinal response to CR we focused on one of the previously observed phenotypes (Duszka et al., 2018), showing increased levels of taurine-conjugated BA in the intestinal contents and are associated with increased levels of free and conjugated taurine in the intestinal mucosa (Figure 4). This observation matches well with elevated expression of BA synthesis- and transport-related genes in the liver of CR mice presented in this study. Correspondingly, the synthesis of taurine is likely to increase in the cysteine dioxygenase-associated cysteine metabolic pathways. However, the expression of the taurine transporter in the liver was not affected, suggesting that taurine is not exported from the liver as free taurine but as bile conjugate. Further, we report that during CR, the levels of taurine increase along the small intestine but not in the liver. Our results show that the amount of BA-derived taurine increases in the CR intestine and various types of conjugates are created to bind taurine. Conjugation of taurine to GSH enhances taurine reuptake from the intestine. Importantly, the impact of GSH on taurine uptake has never been described before and molecular background of this interaction is not known. Correspondingly, we showed (Duszka et al., 2018) that the level of taurine bound to BAs as well as free taurine is lower in the feces of CR compared to *ad libitum* mice indicating increased efficiency of uptake in the small intestine. Accordingly, fasting elevates levels of taurine in urine and/or blood in dogs (Gray et al., 2016), rats (Chesney et al., 1982, Wu, 1954), and pigs (Edmonds and Baker, 1987), with varying effects which depend on the length of fasting.

**Figure 4.**
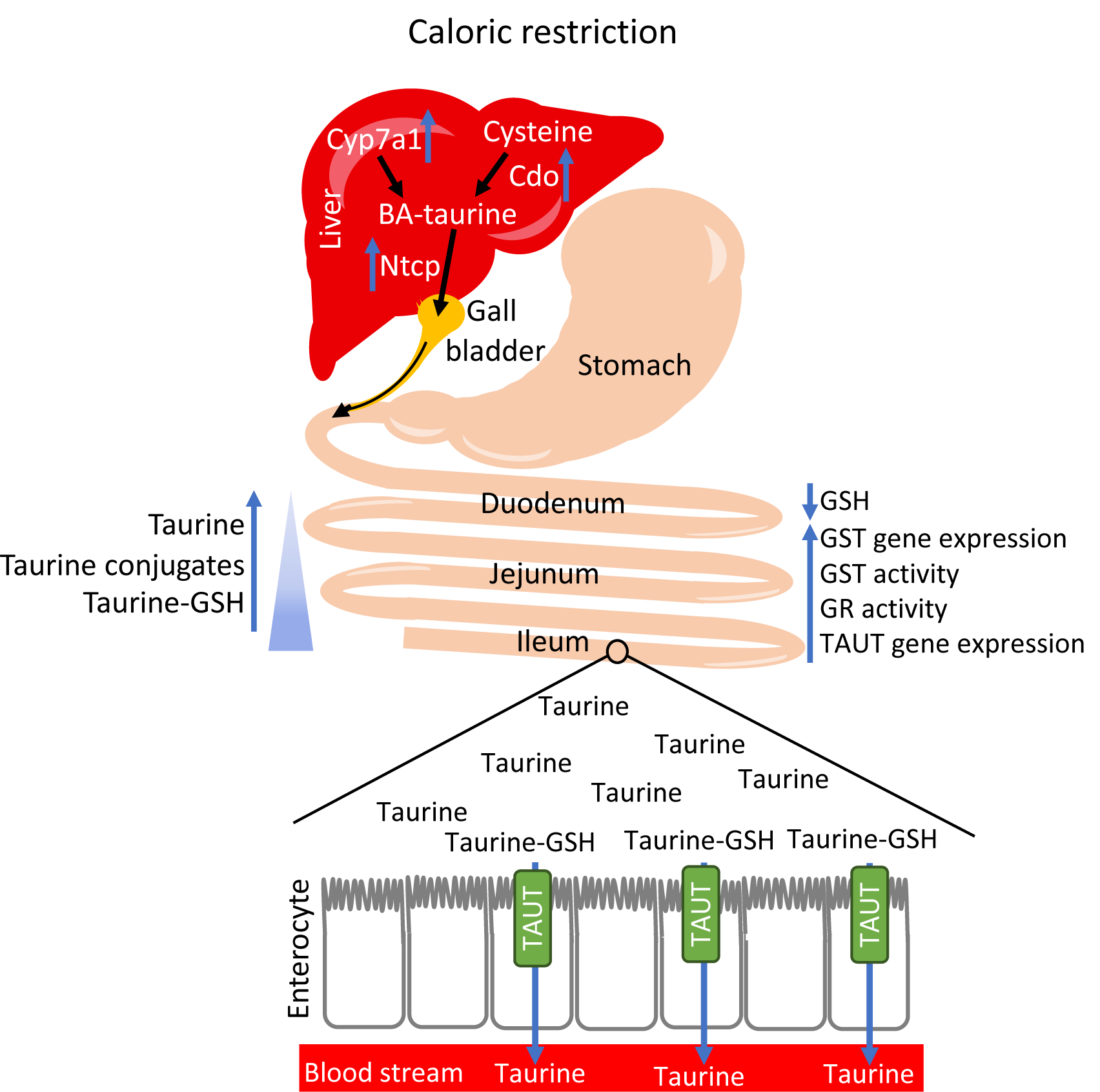
Summary of the results and hypothesis. CR is accompanied by increased expression of genes involved in the production of bile acids (BA) and taurine as well as the secretion of BA from the liver. Taurine-conjugated BA are transported and released into the duodenum. Taurine is deconjugated from BA and creates various types of conjugates which amount increases along the small intestine. The abundance of free taurine stimulates expression and activity of GST leading to the utilization of GSH to create taurine conjugates. Binding GSH results in an imbalance in the free GSH/GSSG ratio which elicits the activity of GR to recreate the homeostasis. CR-triggered increased expression of *TauT* along with conjugation of taurine by GSH enhances taurine uptake from the intestine.

The BAs are secreted to the proximal intestine upon lipid consumption and travel along the GI tract to solubilize lipids aiding digestion and absorption. Once reaching the distal intestine, 95% of BAs are reabsorbed and recirculated to the liver (de Aguiar Vallim et al., 2013). The amount of free taurine, GSH-taurine, and other taurine conjugates are increased in all parts of the intestinal mucosa in CR compared to *ad libitum* mice. However, in this study, the differences between CR and *ad libitum* fed mice are stronger in the distal intestine compared to the duodenum. Most likely, this is due to increased BA reabsorption and high expression of taurine transporter TauT in the ileum (Anderson et al., 2009a).

While, as we showed previously, CR results in the reduction of levels of multiple metabolites in the GI tract (Duszka et al., 2018), taurine is an intriguing example of a molecule with an increased concentration, suggesting a physiological role. The role of enterohepatic circulation or accumulation of taurine during CR has never been investigated. However, we hypothesize that taurine could serve as a regulator of metabolic adjustment as it had been shown to reduce the hepatic secretion of lipids while enhancing fatty acid oxidation and ketogenesis which are processes characteristic to fasting and CR (Fukuda et al., 2011). Additionally, as published before (Duszka et al., 2018), CR results in the downregulation of the immune and anti-microbial genes in the intestine. Fittingly, taurine (Kim and Cha, 2014, Marcinkiewicz et al., 2005, Niu et al., 2018, Chupel et al., 2018, Yu et al., 2016), as well as BAs (Torres et al., 2018, Hang et al., 2019, Song et al., 2020) are known for their anti-inflammatory role and impact on microbiota. Therefore, the increased levels of BAs and taurine in the intestine during CR could contribute to the previously described phenotype. Importantly, taurine supplementation, similary like CR has been shown to reduce endoplasmic reticulum stress and extend lifespan in *C. elegans* (Kim et al., 2010, Wu et al., 2020). Consequently, taurine has already been suggested as a supplement mimicking beneficial outcomes of CR (Nishizono et al., 2017). Seeing the changes of GSH levels in the context of CR we, first, hypothesized changes in the REDOX response, which are often associated with CR. However, contradictory data concerning oxidative stress in CR have been reported (Barja, 2004, Forster et al., 2000, Lambert and Merry, 2005, Lambert et al., 2004, Agarwal et al., 2005). Our results show that intestinal mucosa does not respond to CR by modulating the levels of reactive oxygen species or anti-oxidative capacity. Therefore, we conclude that it is not the oxidative but the conjugative activity of GSH that is involved in the intestinal response to CR and that GST is activated during CR to conjugate taurine which is likely released from BAs. We show that GST gene expression increases and accordingly, the activity of GST and the occurrence of GSH conjugate are elevated in the mucosa of CR mice. Due to the higher demand from the GST side, the level of free GSH decreases. Consequently, the GSSG/GSH ratio is elevated, and this activates GR which converts GSSG to GSH and reestablishes the basic state ratio. It is, however, possible that the decreased level of GSH could also partly result from the CR-accompanied shortage of nutrients, substrates to synthesize GSH, and food-derived GSH. The reduced expression of *Ggt*, coding for GSH hydroxylase which digests external GSH in order to deliver amino acids for absorption and intracellular GSH synthesis, may reflect the reduced availability of the extracellular GSH.

In summary, we describe a novel role of GSH in the handling of CR-associated abundance of taurine in the small intestine. Interestingly, the role of BAs secreted not in the context of food consumption is not known. Importantly, the here described CR-triggered upregulation of GST expression and activity, a link between intestinal GSH and BA-derived taurine as well as the impact of GSH on taurine absorption has never been established before. The phenomenon is likely contributing to the beneficial impact of CR on the GI tract. However, the exact physiological purpose and consequences of this regulation need to be and currently are further investigated.

## Materials and methods

### Animal care and experimental procedures

Male C57Bl/6 mice purchased from Janvier Labs (Le Genest-Saint-Isle, France) were kept under a 12-h light/12-h dark cycle in standard SPF conditions. The animals were fed V153x R/M-H auto diet from SSNIFF-Spezialdiäten GmbH (Soest, Germany) and housed with free water access. Mice aged 12 weeks were randomly divided into experimental groups of nine control *ad libitum* fed (ad lib) or CR mice. During the dissection, some samples were lost resulting in seven to nine biological replicates per the presented data set. The groups did not differ significantly in body weight when starting the experimental procedures. Animal food intake was measured for one week prior to the intervention to determine the amount of chow diet to be given under CR. The mice from the CR group underwent 14 days of CR that consisted of a ∼ 25% reduction of daily food intake. This extent of CR is efficient in triggering CR-related phenotypes but prevents excessive body weight loss as presented previously (Duszka et al., 2018).

All animal experimentation protocols were approved by the Bundesministerium für Wissenschaft, Forschung und Wirtschaft, Referat für Tierversuche und Gentechnik (BMBWF-66.006/0008-V/3b/2018). All experiments were carried out according to animal experimentation Animal Welfare Act guidelines.

### Intestinal sacs assay

The freshly dissected small intestine was divided into five even parts. Part four, counting from duodenum was flushed with PBS and ends were loosely tied with a thread leaving 4 cm-long sac. A blunted needle was introduced and the intestine was filled with 200 μl of taurine (25 mg/ml) or taurine (25 mg/ml) and GSH (61.5 mg/ml) solutions. The sacs were closed tightly and incubated in a 37°C water bath in 10 ml of prewarmed Dulbecco’s Modified Eagle Medium (DMEM). Three 200 μl medium samples were collected over 1.5 h for measurement of taurine transport.

### Protein concentration and activity assays

The levels of GSH and GSSG, as well as the activity of GST, GR, GPx (all from BioVision, CA, USA), and Nrf2 (Abcam, Cambridge, UK), were assessed using commercial assay kits according to the manufacturer’s indications.

### ESR

Frozen samples were cut in 15 μg pieces and 144 μl of oxygen-free KHB and 6μl of oxygen-free 10 mM CMH solution were added. The samples were incubated for 60 min in a 37°C shaking incubator and quickly spun down. 100 μl of the solution from each sample was transferred to a fresh tube and snap-frozen in liquid nitrogen until measuring. ESR measurements were performed at 150 K in a capillary tube (100 μl), which was placed into a high sensitivity resonator (Bruker ER 4122SHQE), using an X-band Bruker Elexsys-II E500 EPR spectrometer (Bruker Biospin GmbH) with a modulation frequency of 100 kHz and a microwave frequency of 9.4 GHz. Spectra were recorded every 20 s, averaging every 10 consecutive spectra. The sweep width was 450 G, the sweep time 20 s, the modulation amplitude 5 G, the center field 3400 G, the microwave power 20 mW, and the resolution was 1024 points. EPR spectra were simulated and the area under the curve determined by double integration of the spectrum. A reference-free quantitation of the number of spins was performed, as has been described previously (Zaunschirm et al., 2018).

### Gene expression

RNA was isolated from intestinal scrapings using the RNeasy mini kit (Qiagen). Samples were thawed in lysis buffer, disrupted using a syringe and needle, and processed following the manufacturer’s recommendations. SuperScript® II Reverse Transcriptase (Invitrogen™, Life Technologies) was used for the reverse transcription step. Quantitative real-time PCR (qRT-PCR) reactions were carried out using the QuantStudio™ 6 Flex Real-Time PCR System (Applied Biosystems, Life Technologies) with the SYBR Green PCR Master Mix (Applied Biosystems, Life Technologies). The primers used are listed in the supplementary data file.

### GSH and taurine conjugates detection

The quantification protocol was adapted from the method of Ito et al. (Ito et al., 2018) and Budinska et al. (Budinska et al., 2020). Frozen liver and intestinal mucosa samples were cut on dry ice to the size of 7-10 mg and homogenized. Liver samples were transferred into Precellys homogenizing tubes with 1.4 mm ceramic beads and nine times the volume of ethanol absolute at −20°C was added. Liver samples were homogenized in the Precellys_®_24 Tissue Homogenizer (Bertin Instruments) twice for 15 s at 5000 rpm, vortexed for 30 s, and incubated at −20°C. Intestinal samples were transferred into 1.5 ml Eppendorf tubes and were disrupted in five thawing and freezing cycles. Next, nine times the volume of ethanol absolute −20°C was added, samples were vortexed for 30 s. After the homogenization, liver and intestinal samples were handled alike. Samples were incubated at −20°C for 24 h and centrifuged for 10 min at 18000 g, the supernatants were transferred to new tubes, and, to remove the remaining debris, the centrifugation step was repeated. The supernatants were transferred into HPLC vials in a thermostatic autosampler kept at 4°C. 60 μl of intestinal sacs samples were diluted with 600 μl ethanol, vortexed mixed, incubated for 20 min at −20°C and centrifuged at 15000 g for 15 min at 4°C. The supernatant was dried in a SpeedVac concentrator for 45 min at 60°C, then dissolved in 70 μl ethanol. Samples (10 μl) were analyzed by LCMS in negative modus using an LCMS-8040 Liquid Chromatograph Mass Spectrometer (Shimadzu Corporation, Kyoto, Japan) with an Atlantis T3 3 μm column (2.1×15 0mm, Waters, Milford, MA, USA). The column temperature was 40°C. The mobile phases consisted of 0.1% formic acid in water (eluent A) and 0.1% formic acid in acetonitrile (eluent B). The gradient was maintained at an initial 5% B for 2.5 min, to 20% B at 8 min, and was set back to 5% B at 9 min with a hold for one minute.

### Identification of conjugates

Standards of GSH and taurine (both from Sigma-Aldrich, St. Louis, MO, USA) were prepared in 70% ethanol. To induce the conjugation of taurine to GSH 150 mmol of each was weighted in the same 2 ml Eppendorf tube, 70% ethanol was added, vortexed shortly, and was incubated for 30 min at room temperature. Standards were separated with the column and HPLC gradient described above and were fragmented with an LC-MS system (LCMS-80-40, Shimadzu, Korneuburg, Austria). The MS instrument was operated in multiple reaction mode (MRM) with the following settings: nebulizing gas flow 3L/min, drying gas flow 12L/min, desolvation line temperature 250°C and heat block temperature 350°C. Argon was used as the collision-induced dissociation (CID) gas with a collision energy of 20eV. The fragmentation pattern was compared to METLIN’s database. To identify GSH and taurine conjugates, samples were screened for precursor ions producing similar fragmentation patterns to GSH and taurine. The exact mass of the selected precursor ions was measured using an LC-ESI-TOF-system consisting of an Ultimate 3000 (Thermo Fischer Scientific, Waltham, Massachusetts, US) and a micrOTOF-Q II (Bruker Daltonics, Bremen, Germany) with an Atlantis T3 3 μm column (2.1×150 mm, Waters, Milford, MA, USA). The column temperature was 40°C. The mobile phases consisted of 0.1% formic acid in water (eluent A) and 0.1% formic acid in acetonitrile (eluent B). The gradient was maintained at an initial 5% B for 2.5 min, to 20% B at 8 min, and was set back to 5% B at 9 min with a hold for one minute. The exact mass and fragmentation pattern of the selected precursor ions were checked against the METLIN database. The compounds identified as GSH or taurine conjugates are listed in the supplementary table 2.

### Statistics

Within the study, data sets with two groups (CR and *ad libitum*) of seven to nine biological replicates were compared using a two-sided student’s t-test to verify statistical significance. A p-value lower than 0.05 was considered statistically significant. The presented data stems from two technical replicates.

## Acknowledgments

The authors would like to thank Dr. Barbara Lieder for her input in the discussions concerning the publication.

Open access funding provided by the University of Vienna.

---

**Supplementary figure 1.**
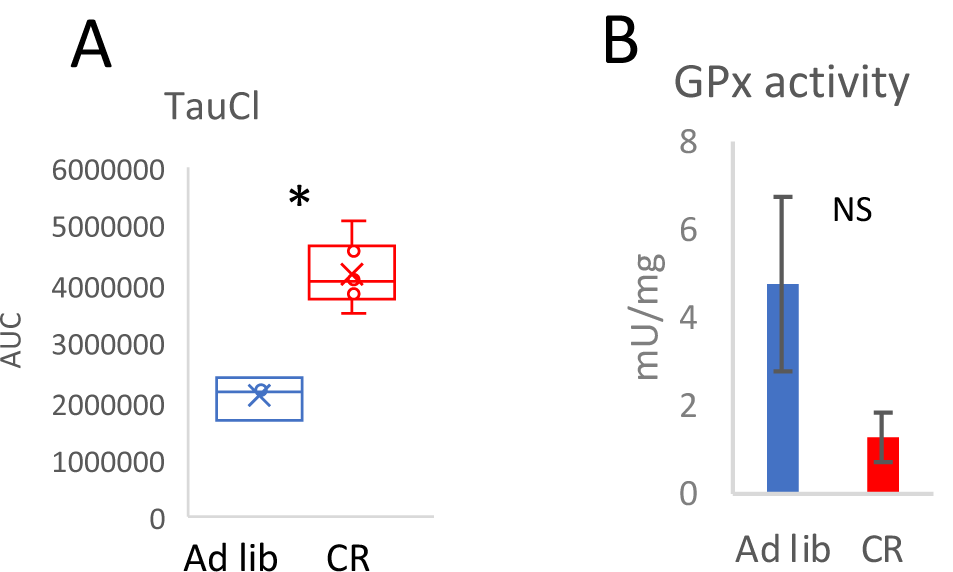

