## Supplementary material for "Caloric restriction increases levels of taurine in the intestine and stimulates taurine uptake by conjugation to glutathione": Supplementary information.pdf

**Supplementary tables**

**Supplementary table 1: Primers list**

| Gene ID | Forward | Reverse |
|---|---|---|
| Eef1a1 | CCTGGCAAGCCCATGTGT | TCATGTCACGAACAGCAAAGC |
| Gclc | ATGTGGACACCCGATGCAGTATT | TGTCTTGCTTGTAGTCAGGATGGTTT |
| Ggt | ACCACTCACCCAACCGCCTAC | ATCCGAACCTTGCCGTCCTT |
| GPx1 | CCCCACTGCGCTCATGA | GGCACACCGGAGACCAAA |
| GPx4 | GCTGTGCGCGCTCCAT | CCATGTGCCCGTCGATGT |
| Grx1 | TGCAGAAAGACCCAAGAAATCCTCAGTCA | TGGAGATTAGATCACTGCATCCGCCTATG |
| Grx2 | CATCCTGCTCTTACTGTTCCATGGCCAA | TCATCTTGTGAAGCGCATCTTGAAACTGG |
| Gs | TCTAACAAGAAACATCCGGCA | GGTCAGGTCGATGTCATTGTA |
| Gsta3 | GAATGGAGCCTATCCGGTGG | GCATGGCGGTACAAGCCTTT |
| Gsta4 | ACTTTAATGGCAGGGGACGG | GCAGGTGTCCATCCTTTTGC |
| Mgst1 | CCTTCTCCCTGGATTCAGTCAT | TCGGCCATGCTTCCAATCTT |
| Mgst2 | GAAAGAAAGATGGCCGGGGA | CCGCCAAGCGAAATAACTTTG |
| Nrf2 | TCTCCTCGCTGGAAAAAGAA | AATGTGCTGGCTGTGCTTTA |

**Supplementary table 2: Detected compounds**

| Precursor ions | Product ions | Retention time (min) | Name of the compound | Tissue of detection |
|---|---|---|---|---|
| 124.0085 | 80 ($SO3^-$) | 2.8 | Taurine | Liver, small intestine |
| 158.8468 | 124 ($C2H6NO3S^-$),<br>80 ($SO3^-$) | 2.7 | Taurine chloramine | Small intestine |
| 249.0212 | 80 ($SO3^-$),<br>124 ($C2H6NO3S^-$) | 2.6 | Unknown taurine conjugate | Liver, small intestine |
| 304.9204 | 80 ($SO3^-$),<br>124 ($C2H6NO3S^-$) | 2.7 | Unknown taurine conjugate | Liver, small intestine |
| 306.0750 | 128 ($C5H6NO3^-$),<br>143 ($C5H7N2O3^-$),<br>160 ($C5H3NO3S^-$),<br>179 ($C8H7N2O3^-$),<br>210 ($C9H12N3O3^-$), | 2.8 | GSH | Liver, small intestine |

| | | | | |
|---|---|---|---|---|
| | 254 (C10H12N3O5-),<br>272 (C10H14N3O6-) | | | |
| 308.0733 | 80 (SO3-),<br>124 (C2H6NO3S-) | 2.6 | Unknown taurine conjugate | Liver, small intestine |
| 404.0561 | 128 (C5H6NO3-),<br>143 (C5H7N2O3-),<br>160 (C5H3NO3S-),<br>179 (C8H7N2O3-),<br>210 (C9H12N3O3-),<br>254 (C10H12N3O5-),<br>272 (C10H14N3O6-) | 2.9 | Unknown GSH conjugate | Liver, small intestine |
| 431.0943 | 80 (SO3-),<br>124 (C2H6NO3S-),<br>128 (C5H6NO3-),<br>143 (C5H7N2O3-),<br>160 (C5H3NO3S-),<br>179 (C8H7N2O3-),<br>210 (C9H12N3O3-),<br>254 (C10H12N3O5-),<br>272 (C10H14N3O6-) | 2.7 | GSH-taurine conjugate | Liver, small intestine |

**Supplementary figure legend**

**Supplementary figure 1**

The levels of Taurine chloramine (TauCl) in the jejunum (A). Glutathione peroxidase (GPx) activity was measured using a commercial assay (B). Two-tailed Student's t-test was applied to assess statistical differences between the groups; *$p<0.05$. The bars indicate the mean of eight to nine biological replicates ±SEM.

Supplementary figure 1

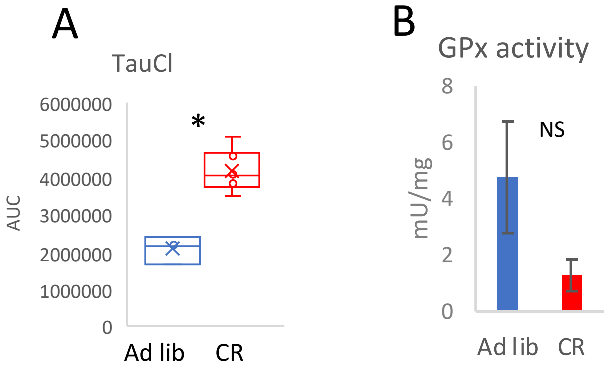
